# Cyclic oligoadenylate signalling mediates *Mycobacterium tuberculosis* CRISPR defence

**DOI:** 10.1101/667758

**Authors:** Sabine Grüschow, Januka S. Athukoralage, Shirley Graham, Tess Hoogeboom, Malcolm F. White

## Abstract

The CRISPR system provides adaptive immunity against mobile genetic elements (MGE) in prokaryotes. In type III CRISPR systems, an effector complex programmed by CRISPR RNA detects invading RNA, triggering a multi-layered defence that includes target RNA cleavage, licencing of an HD DNA nuclease domain and synthesis of cyclic oligoadenylate (cOA) molecules. cOA activates the Csx1/Csm6 family of effectors, which degrade RNA non-specifically to enhance immunity. Type III systems are found in diverse archaea and bacteria, including the human pathogen *Mycobacterium tuberculosis*. Here, we report a comprehensive analysis of the *in vitro* and *in vivo* activities of the type III-A *M. tuberculosis* CRISPR system. We demonstrate that immunity against MGE is achieved predominantly via a cyclic hexa-adenylate (cA_6_) signalling pathway and the ribonuclease Csm6, rather than through DNA cleavage by the HD domain. Furthermore, we show that the mechanism can be reprogrammed to operate as a cyclic tetra-adenylate (cA_4_) system by replacing the effector protein. These observations demonstrate that *M. tuberculosis* has a fully-functioning CRISPR interference system that generates a range of cyclic and linear oligonucleotides of known and unknown functions, potentiating fundamental and applied studies.

## INTRODUCTION

The CRISPR system provides adaptive immunity against Mobile Genetic Elements (MGE) in prokaryotes (reviewed in (1,2)). CRISPR systems are classified into two major classes and six major types: class 1 (types I, III and IV) have multi-subunit effector complexes based around a Cas7 backbone subunit, whilst class 2 (types II, V and VI) are single-subunit enzymes (3). The latter, encompassing Cas9, Cas12 and Cas13, have shown great utility in genome engineering applications due to their relative simplicity. However, class 1 systems are more widespread in prokaryotes. Type III effector systems are arguably the most complex, powerful and versatile of the characterised CRISPR interference complexes. Assembled around a Cas7 backbone that binds CRISPR RNA (crRNA) (4), they act as surveillance complexes in the cell, detecting target RNA molecules from MGE via base-pairing to the crRNA. Target RNA binding licenses the catalytic Cas10 subunit to degrade DNA, via its HD-nuclease domain (5-10) and to synthesise cyclic oligoadenylate (cOA) molecules via its Cyclase/Palm domain (11-13). cOA, which is assembled from 3-6 AMP monomers to generate species from cA_3_ to cA_6_, is a potent second messenger that in turn binds to downstream effector complexes via a CRISPR Associated Rossman Fold (CARF) domain (14). Diverse proteins with CARF domains have been identified bioinformatically, but only one family has been studied extensively: the Csx1/Csm6 family that utilises a C-terminal HEPN (Higher Eukaryotes and Prokaryotes, Nucleotide binding) domain to cleave RNA (15,16). These enzymes are important for CRISPR-based immunity *in vivo* (17-19). Thus, type III CRISPR systems are capable of directing a multi-faceted antiviral defence on detection of foreign RNA in the cell (Figure 1A).

**Figure 1.**
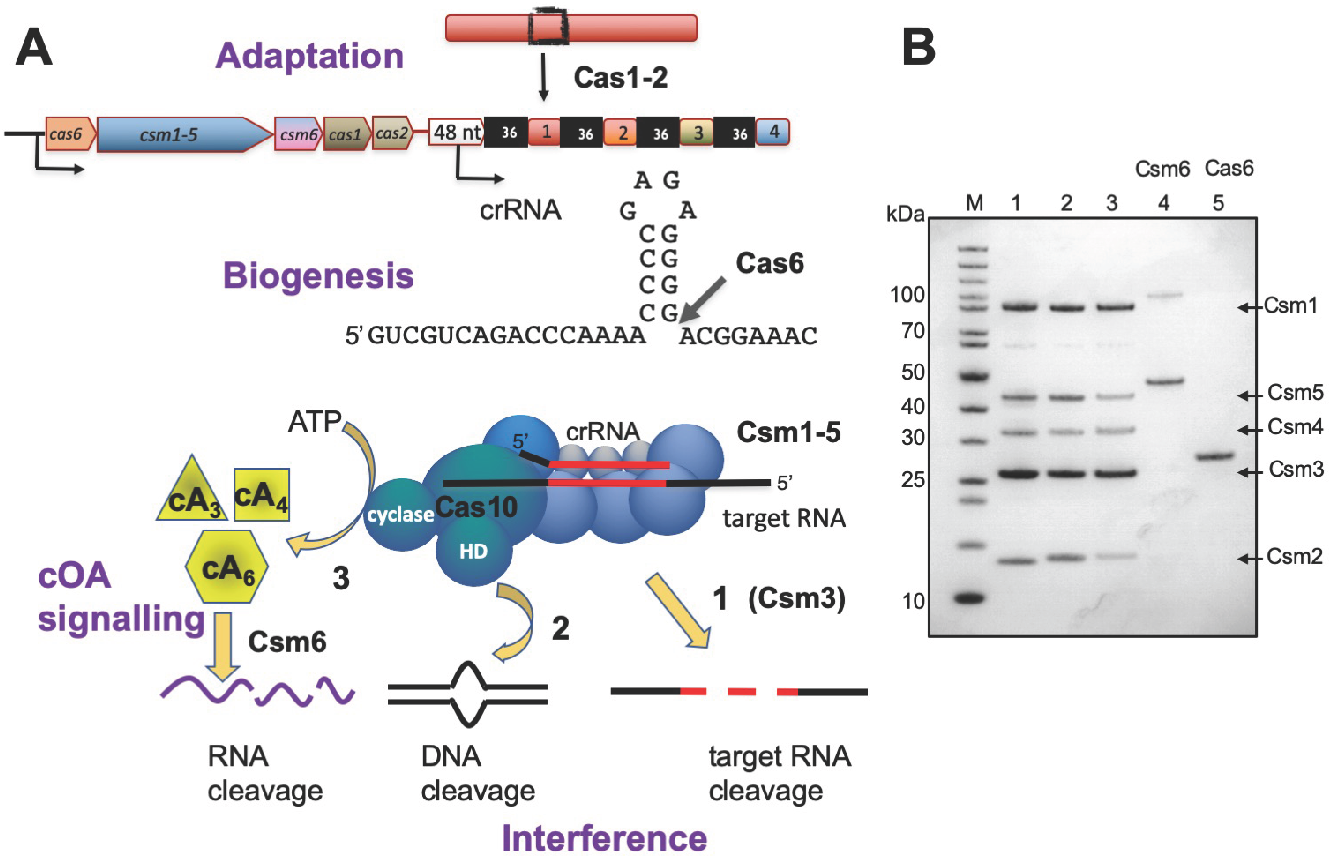
The CRISPR system of *M. tuberculosis*. A. The CRISPR locus of *M. tuberculosis* includes genes encoding Cas6 (crRNA processing), Csm1-5 (type III-A interference complex), Csm6 (ancillary ribonuclease), Cas1 and Cas2 (Adaptation). Cas6 cleaves the CRISPR RNA at the base of a short hairpin to generate mature crRNA that is bound by the Csm complex. On target RNA binding, the Csm complex is expected to display three enzymatic activities: target RNA cleavage (1), DNA cleavage by the HD domain (2) and cOA production by the cyclase domain (3). B. Purified, recombinant CRISPR-associated proteins of *M. tuberculosis*. M: PageRuler Unstained (Thermo Scientific); 1: Csm1-5 interference complex; 2: Csm1-5, Csm1 D630A, D631A (Cy variant); 3: Csm1-5, Csm3 D35A (C3 variant); 4: Csm6; 5: Cas6.

*Mycobacterium tuberculosis* (*Mtb*) is the causative agent of tuberculosis (TB), which imposes a huge burden on human health and on developing economies, with 10 million new infections and 1.6 million deaths in 2017 (WHO Global Tuberculosis Report, 2018). The majority of *Mtb* strains encode a complete CRISPR system (20), and indeed the type III-A (or Csm) CRISPR system was originally labelled as the “Mtube” subtype based on the genes present in *Mtb* strain H37Rv (21). CRISPR loci vary between *Mtb* strains and have been used for many years as a strain typing (spoligotyping) tool (22), suggesting uptake of DNA derived from mycobacteriophage or other MGE by the CRISPR system occurs. The type III-A system encoded by *Mtb* H37Rv comprises Csm1-6, with Cas6 for crRNA processing and the adaptation proteins Cas1 and Cas2 next to the CRISPR array (Figure 1A). In *M. bovis* (BCG), the organisation is identical with the exception of a truncating mutation in the *csm6* gene (20). Looking across the entirety of the prokaryotic domains of life, *Mtb* is a rare example of an isolated type III effector and associated adaptation system.

Little is known about the activity of the *Mtb* CRISPR system, although recent work suggests it can provide modest immunity against MGE (23). Here, we report the reconstitution and characterisation of the *Mtb* system both *in vitro* and *in vivo*, in *E. coli*. We demonstrate that the *Mtb* Csm complex provides immunity via a cA_6_-activated Csm6 ribonuclease, and that the system can be reprogrammed to provide cA_4_-modulated immunity by swapping the cOA-activated effector protein.

## MATERIALS AND METHODS

### Cloning

Enzymes were purchased from Thermo Scientific, BioLine, Promega or New England Biolabs and used according to manufacturer’s instructions. Oligonucleotides (Table S2) and synthetic genes (Table S3) were obtained from Integrated DNA Technologies (Coralville, Iowa, USA). Synthetic genes were codon-optimized for *E. coli* and restriction sites for cloning incorporated. All constructs were verified by sequencing (GATC Biotech, Eurofins Genomics, DE) unless stated otherwise. Protein concentrations were determined by UV quantitation (NanoDrop 2000, Thermo Scientific) using calculated extinction coefficients (ExPASy, ProtParam software) unless stated otherwise.

#### CRISPR genes

All genes were obtained as synthetic genes. To obtain N-terminal His_6_-fusion proteins, *Mtb cas10* (*csm1*), *csm2, csm3, csm4, csm5*, and *csm6* were digested with *Nco*I and *Xho*I and ligated into the corresponding sites of the expression vector pEHisTEV (24). The *Mtb cas6* gene was cloned into the *Nde*I and *Xho*I sites of the same vector to obtain the C-terminal His_6_-fusion. We cloned a truncated version of *cas6*, starting at Met53 relative to NCBI annotation, because the N-terminal region showed no homology to non-mycobacterial Cas6 proteins, and a transcription start site was identified within the predicted *cas6* gene (25). *Thioalkalivibrio sulfidiphilus csx1* was cloned into the *Nco*I and *Bam*HI sites of pEV5HisTEV to obtain a N-terminal His_8_-fusion (13).

#### CRISPR arrays

A CRISPR array containing three *Mtb* repeat sequences flanking two identical spacer sequences targeting the pUC19 multiple cloning site was assembled by *Bam*HI digest of the double-stranded mtbCRISPR array oligonucleotide (Table S2) followed by ligation into pCDFDuet™-1 (Novagen, Merck Millipore). *Mtb cas6* was cloned into MCS-2 to give pCRISPR.

A CRISPR array consisting of four identical spacers targeting the tetracycline resistance gene, flanked by five *Mtb* repeat sequences, was assembled from phosphorylated oligonucleotides mtbCA_Rep and mtbCA_TetR-Sp (Table S2). The array was ligated into MCS-1 of pCDFDuet™-1 containing *cas6* in MCS-2 to give pCRISPR-Target.

#### *Mtb* Csm complex for *in vitro* studies

*Cas10* (*csm1*), *csm2*, and *csm5* were cloned into pACYCDuet-1 (5’-*Nco*I, 3’-*Xho*I; Novagen, Merck Millipore), *csm4* was cloned into pEHisTEV (5’-*Nco*I, 3’-*Xho*I) and *csm3* was inserted into pEHisTEV (5’-NdeI, 3’-XhoI) by restriction enzyme cloning. These constructs were used as templates for assembly of *M. tuberculosis* CRISPR genes, including ribosome binding sites, into MultiColi™ (Geneva Biotech, Genève, CH) acceptor and donor vectors by sequence- and ligation-independent cloning (SLIC) as per manufacturer’s instructions. Briefly, *His*_*6*_*-csm4, csm2*, and *cas10* were introduced into the acceptor pACE; *csm3, csm5* were cloned into pDK; *Mca csm2* was inserted into pDC using restriction enzyme cloning (5’-*Nde*I, 3’-*Xho*I). The acceptor and donor plasmids were then recombined using Cre recombinase to give pCsm1-5. To obtain variant Csm complexes, His_8_-V5-*csm4* (from pEHisV5TEV-Csm4), *Mca csm2* (from pACYCDuet-Mca_Csm2) and *cas10* (*csm1*) were introduced into pACE by SLIC; and *csm3, csm5* were inserted into the donor plasmid pDK. Primer-directed mutagenesis was carried out on the assembled acceptor and donor plasmids, before Cre recombination to give pCsm1-5_C3 (Csm3 D35A) and pCsm1-5_Cy (Csm1 D630A/D631A). The identity of all donor and acceptor plasmids was verified by sequencing, and the final, recombined construct was analysed by restriction digest.

#### *Mtb* Csm complex for *in vivo* studies

The same templates as for construction of pCsm1-5_C3 were used with the addition of *csm6* (pACYCDuet-1). The pACE assembly was as for pCsm1-5_C3. *Csm3, csm5*, and *csm6* were introduced into pDK; for the ΔCsm6 plasmid, the donor plasmid *en route* to pCsm1-5 was used. Mutagenesis was carried out as before. All constructs were verified by sequencing. Suitable acceptor and donor plasmid combinations were then assembled by Cre recombination to give pCsm1-6, pCsm1-5_ΔCsm6, pCsm1-6_C3, pCsm1-6_Cy, and pCsm1-6_HD (Table S1). The control plasmid pCsm_Control was obtained by Cre recombination of pACE and pDK without inserts. The identity of the final, recombined constructs was confirmed by restriction digest analysis.

#### Construction of target plasmids

An arabinose-inducible pRSFDuet™-1 (Novagen, Merck Millipore) derivative with tetracycline resistance was constructed as follows. The *araC* and pBAD region was PCR-amplified from pBADHisTEV (Huanting Liu, University of St Andrews, UK) using primer pair ara-*Avr*II-R/F and cloned into the *Xba*I and *Nde*I sites of pRSFDuet™-1 to give pRSFara. The tetracycline resistance gene was PCR-amplified from the MultiColi™ vector pACE2 using the TetR primer pair; the pRSFara-backbone lacking its resistance gene was PCR-amplified using primer pair RSFara. The two PCR products were digested with *Sph*I and *Bam*HI and joined through ligation to give pRAT. The pUC19 MCS / *lacZα* gene was introduced into pRAT by megaprimer PCR (primer pair lacZ-pRAT) to give pRAT-Target (Table S1).

### Protein Production and Purification

#### *Mtb* Cas6

C-terminally His_6_-tagged Cas6 was expressed in BL21 (DE3) grown in LB containing 50 µg ml^-1^ kanamycin and 50 µg ml^-1^ chloramphenicol. An overnight culture was diluted 100-fold into fresh medium and grown at 37 °C, 180 rpm to an OD_600_ of 0.4. The temperature was lowered to 16 °C, and incubation continued until the OD_600_ reached 0.8. Production was induced with 502 µM IPTG, and the cultures were continued for a further 20 h. Cells were harvested by centrifugation. The pellet was resuspended in 50 mM Tris-HCl, 250 mM NaCl, 10 mM imidazole, 10 % glycerol, pH 7.5, sonicated, and the cell debris removed by centrifugation. The cleared lysate was loaded onto a pre-equilibrated HisTrap Crude FF (GE Healthcare) column, washed with 50 mM Tris-HCl, 250 mM NaCl, 30 mM imidazole, 10 % glycerol, pH 7.5 and eluted in a gradient to 50 mM Tris, 250 mM NaCl, 500 mM imidazole, 10 % glycerol, pH 7.5. Cas6-containing fractions were pooled, and further purified by gel filtration (HiLoad 16/60 Superdex pg 200, GE Healthcare) in 50 mM Tris-HCl, 250 mM NaCl, 10 % glycerol, pH 7.5. Cas6 and Cas6 H47N were concentrated as required using an Amicon Ultra centrifugal filter (MWCO 10 kDa, Merck-Millipore).

#### *Mtb* Csm complex

*E. coli* C43(DE3) cells were co-transformed with the MultiColi™ construct pCsm1-5 (or pCsm1-5_C3, pCsm1-5_Cy) and pCRISPR. Overnight cultures were diluted 100-fold into LB containing 100 µg ml^-1^ ampicillin and 50 µg ml^-1^ spectinomycin, incubated at 37 °C, 180 rpm until the OD_600_ reached 0.8. After induction with 100 µM IPTG, incubation was continued at 25 °C for 5 h. Cells were harvested by centrifugation and pellets stored at −20 °C. Cells were resuspended in lysis buffer (50 mM Tris-HCl, 500 mM NaCl, 10 mM imidazole, 10 % glycerol, pH 7.5) and lysed by sonication. The cleared lysate was loaded onto HisPur Ni-NTA resin (Thermo Scientific), washed with lysis buffer and eluted with lysis buffer containing 250 mM imidazole. Csm complex-containing fractions were diluted 2-fold with lysis buffer to prevent aggregation, pooled, then dialysed at 4 °C overnight in the presence of 0.3 mg ml^-1^ TEV protease against lysis buffer. The protein solution was passed through His-Pur Ni-NTA resin a second time, and the flow-through was concentrated using an Amicon Ultracentrifugal filter (30 kDa MWCO, Merck-Millipore) and further purified by gel filtration (HiPrep 16/60 Sephacryl pg 300 HR, GE Healthcare) using 20 mM Tris-HCl, 250 mM NaCl, 10 % glycerol, pH 7.5 as mobile phase. When necessary, the Csm complex was further purified on MonoS 4.6/100 PE (GE Healthcare) in 20 mM MES, 50 mM NaCl, pH 6 and eluted using a salt gradient to 1 M NaCl. Single-use aliquots were flash-frozen and stored at −80 °C.

#### *Mtb* Csm6 and *T. sulfidiphilus* Csx1

*Mtb* Csm6 and TsuCsx1 were expressed in C43 cells. Cells were grown for 4 h at 25 °C before harvest by centrifugation at 4000 rpm at 4 °C for 15 min (JLA 8.1 rotor). Cells were lysed in buffer containing 50 mM Tris-HCl pH 8.0, 0.5 M NaCl, 10 mM imidazole and 10% glycerol supplemented with 1 mg/ml chicken egg lysozyme (Sigma-Aldrich) and one EDTA-free protease inhibitor tablet (Roche) by sonicating six times for 1 min with 1 min rest intervals on ice. Cell lysate was then ultracentrifugated at 40000 rpm (70 Ti rotor) at 4 °C for 45 min and loaded onto a 5 ml HisTrap FF Crude column equilibrated with wash buffer containing 50 mM Tris-HCl pH 8.0, 0.5 M NaCl, 30 mM imidazole and 10 % glycerol. After washing unbound protein with 20 CV wash buffer, recombinant Csm6 or TsuCsx1 was eluted with a linear gradient of wash buffer supplemented with 0.5 M imidazole across 16 CV, holding at 10% for 2 CV, 20% for 4 CV and 50% for 3 CV. Csm6 could not be concentrated and was therefore further purified using MonoQ (GE Healthcare) anion exchange chromatography, diluting protein from nickel affinity fractions directly in buffer containing 20 mM Tris-HCl pH 7.5 and 50 mM NaCl and eluting protein across a linear gradient of buffer containing 20 mM Tris-HCl pH 7.5 and 1M NaCl. For TsuCsx1, nickel affinity fractions containing the protein were concentrated using a 10 kDa MWCO ultracentrifugal concentrator, and further purified by size exclusion chromatography (S200 26/60; GE Healthcare) in buffer containing 20 mM Tris-HCl pH 8.0, 0.5 M NaCl and 1 mM DTT. All proteins were aliquoted, flash frozen with liquid nitrogen, and stored at −80 °C.

### Assays

#### Cyclic oligonucleotides

cA_4_ and cA_6_ were purchased from BIOLOG Life Science Institute (Bremen, DE).

#### Oligonucleotides

All RNA and DNA oligonucleotides as well as 6-FAM™-labeled RNA substrates were purchased from Integrated DNA Technologies (Leuven, BE). For nuclease assays, labelled oligonucleotides were gel purified as described previously (26). Duplexed DNA oligonucleotides were prepared by mixing equimolar amounts of ssDNA in 10 mM Tris-HCl, 50 mM NaCl, 0.5 mM EDTA, pH 8.0, heating to 95 °C for 10 min, followed by slow cooling. Oligonucleotides were 5’-end-labeled with [γ-^32^P]-ATP (10 mCi ml^-1^, 3000 Ci mmol^-1^; Perkin Elmer) using polynucleotide kinase (Thermo Scientific). Oligonucleotides were quantified spectophotometrically using calculated extinction coefficients (OligoAnalyzer tool, IDT). RNA ladders were obtained by alkaline hydrolysis (Thermo Fisher Scientific, RNA Protocols).

#### Cas6 ribonuclease assay

The reaction mixture contained 3 μM Cas6 in 20 mM Tris-HCl, 100 mM potassium glutamate, 1 mM DTT, 5 mM EDTA, 0.1 U µl^-1^ SUPERase•In™ (Thermo Scientific), pH 7.5. Reactions were initiated by addition of 5’ 6-FAM™-labeled mtbRepeat_Cas6 RNA (Table S2) to 50 nM final concentration. The samples were incubated for up to 1 h at 37 °C. The reaction was stopped by extraction with phenol-chloroform-isoamyl alcohol. RNA species were resolved on a 20 % denaturing (7 M urea) acrylamide gel in 1X TBE buffer, and visualised by scanning (Typhoon FLA 7000, GE Healthcare).

#### Target RNA cleavage by the Csm complex

The ribonuclease activity of the Csm complex was assessed by adding 25 nM [5’-^32^P]-labeled target RNA (Table S2) to 0.8 µM Csm, 5 mM MgCl_2_, 0.1 U µl^-1^ SUPERase•In™ (Thermo Scientific), 50 mM Tris-HCl, 50 mM NaCl, pH 8.0, and incubating at 30 °C for up to 2 h. The reaction was stopped by phenol-chloroform-isoamyl alcohol extraction to remove proteins. P1.A26 RNA (Table S2) was used as non-target RNA control. Products were separated by gel electrophoresis (20 % denaturing polyacrylamide) and visualised by exposure to a BAS Storage Phosphor Screen (GE Healthcare).

#### DNase activity of the Csm complex

To test for DNase activity of the Csm complex, 25 nM 5’-_32_P-labeled DNA substrate was added to 0.8 µM Csm, 200 nM3 target RNA, 0.1 mM CoCl_2_, 0.1 U µl^-1^ SUPERase•In™ (Thermo Scientific), 50 mM Tris-HCl, 50 mM NaCl, pH 8.0. The solution was incubated at 30 °C for 90 min, and the reaction stopped by phenol-chloroform extraction. Products were analysed as described above.

#### Cyclic oligoadenylate production and LC-MS analysis

Cyclic oligoadenylate production was triggered by adding 2 µM crude target RNA to 0.8 µM Csm, 1 mM ATP, 5 mM MgCl_2_, 0.1 U µl^-1^ SUPERase•In™ (Thermo Scientific), 50 mM Tris-HCl, 50 mM NaCl, pH 8.0, and incubating at 30 °C for up to 2 h. The reaction was stoped by phenol-chloroform-isoamyl alcohol extraction. Reaction mixtures that were used for Csm6 or Tsu Csx1 activation assays were further extracted with chloroform-isoamyl alcohol to remove residual phenol. Liquid chromatography-high resolution mass spectrometry (LC-HRMS) analysis was performed on a Thermo Scientific Velos Pro instrument equipped with HESI source and Dionex UltiMate 3000 chromatography system. Samples were desalted on C18 cartridges (Harvard Apparatus). Compounds were separated on a Kinetex 2.6 µm EVO C18 column (2.1 × 5 mm, Phenomenex) using the following gradient of acetonitrile (B) against 20 mM ammonium bicarbonate (A): 0 – 2 min 2 % B, 2 – 10 min 2 – 8 % B, 10 – 11 min 8 – 95 % B, 11 – 14 min 95 % B, 14 – 15 min 95 – 2 % B, 15 – 20 min 2 % B at a flow rate of 350 µl min^-1^ and column temperature of 40 °C. UV data were recorded at 254 nm. Mass data were acquired on the FT mass analyzer in positive ion mode with scan range *m/z* 200 – 2000 at a resolution of 30,000.

#### Csm6 activation assay

Csm6 (125 nM dimer) was incubated with 25 nM 5’-radiolabeled P1.A26 RNA substrate together with 2 µM cold RNA in 50 mM MES, 100 mM potassium L-glutamate, pH 6.5. The reaction was started by addition of synthetic cA_6_ or 0.1 volumes protein-free reaction mixture from the cyclic oligoadenylate production above. The reaction was stopped after 45 min at 37 ° C by phenol-chloroform extraction. Products were analysed by 20 % denaturing polyacrylamide gel electrophoresis.

#### TsuCsx1 activation assay

Csx1 (500 nM dimer) was incubated with 50 nM 5’-radiolabeled A1 RNA substrate and 1 µM cA_4_, 1 µM cA_6_ or 1 µl Csm-derived cOA (see above) in pH 7.5 buffer containing 20 mM Tris-HCl pH 7.5, 150 mM NaCl and 1 mM DTT in a 10 µl reaction volume at 35 °C. Reactions were stopped at 1, 5, 15 or 30 minutes by the addition of a reaction volume equivalent of phenol-chloroform and extracted products were analysed by denaturing gel electrophoresis as described above.

#### Plasmid immunity *in vivo*

pCsm1-6 and pCRISPR were co-transformed into *E. coli* C43(DE3). Plasmids were maintained by selection with 100 µg ml^-1^ ampicillin and 50 g ml^-1^ spectinomycin. Competent cells were prepared by diluting an overnight culture 50-fold into fresh, selective LB medium; the culture was incubated at 37 °C, 220 rpm until the OD_600_ reached 0.8 – 1.0. Cells were collected by centrifugation and the pellet resuspended in an equal volume of 60 mM CaCl_2_, 25 mM MES, pH 5.8, 5 mM MgCl_2_, 5 mM MnCl_2_. Following incubation on ice for 1 h, cells were collected and resuspended in 0.1 volumes of the same buffer containing 10 % glycerol. Aliquots (100 µl) were stored at −80 °C. Target or control plasmid (100 ng pRAT-Target, 100 ng pRAT, respectively) were added to the competent cells, incubated on ice for 30 min and transformed by heat shock. LB medium (0.5 ml) was added and the transformation mixture incubated with shaking for 2.5 h before collecting the cells and resuspending in 100 µl LB. 5 µl of a 10-fold serial dilution were applied in duplicate to LB agar plates supplemented with 100 µg ml^-1^ ampicillin and 50 g ml^-1^ spectinomycin when selecting for recipients only; transformants were selected on LB agar containing 100 µg ml^-1^ ampicillin, 50 g ml^-1^ spectinomycin, 25 µg ml^-1^ tetracycline, 0.2 % (*w/v*) β-lactose and 0.2 % (*w/v*) L-arabinose. Arabinose was omitted when transcriptional induction of target was not required. Plates were incubated at 37 °C for 16 – 18 h. Colonies were counted manually and corrected for dilution and volume to obtain colony-forming units (cfu) ml^-1^, statistical analysis was performed with RStudio (RStudio Inc., Boston, USA, version 1.2.1335). The experiment was performed as two independent experiments with two biological replicates and at least two technical replicates.

## RESULTS

### Production of the functional Mtb Csm complex

Although the *Mtb* type III-A effector complex was described very early in the classification of CRISPR systems (21), there have been no subsequent publications describing studies of the recombinant system. This is surprising given the importance of *Mtb* as a human pathogen and the relative lack of model type III CRISPR systems outwith the hyperthermophiles. We took the initial approach of expressing synthetic genes encoding Csm1-5 individually in E. coli using the expression vector pEhisV5Tev (27). Most of the subunits were expressed strongly, although several had solubility problems. We also expressed the *Mtb* Csm6 and Cas6 enzymes in soluble form (Figure 1B). Cas6 was observed to cleave a CRISPR RNA substrate to generate the expected 8 nt 5’-handle (Figure S1). This activity was stimulated by the addition of a divalent cation, as observed previously (23). However, the ability of Ca^2+^ to support catalysis is suggestive of a role for the metal ion in RNA folding and stability rather than a direct role in the catalytic mechanism, and all other Cas6 enzymes studied are metal-independent enzymes (reviewed in (28)).

We next constructed a plasmid to express all five subunits of *Mtb* Csm simultaneously using the MultiColi™ system (Geneva Biotech), along with a compatible pCDFDuet-1 derivative expressing an *Mtb* mini-CRISPR and the Cas6 enzyme (pCRISPR, Table S1) to allow for co-production of the crRNA-charged interference complex in *E. coli*. Using a polyhistidine tag on the Csm4 subunit, it was noted that all subunits except Csm2 could be identified. We therefore added a gene encoding the Csm2 homologue from *Mycobacterium canettii* (*Mca*) to the Csm expression vector (pCsm1-5, Table S1). The two proteins are 71 % identical at the amino acid level. Gratifyingly, the Csm complex could then be purified using affinity chromatography followed by gel filtration (Figure 1B). Mass spectrometry confirmed that all subunits were present. In all subsequent Csm expression plasmids, the *Mtb csm2* gene was therefore replaced by *Mca csm2*. Protein expression plasmids were also constructed carrying site directed mutations targeted to the active sites of the Csm complex: a Cyclase (Cy) variant (GGDD to GGAA, pCsm1-5_Cy) and a Csm3 D35A variant (C3, pCsm1-5_C3, Table S1), as described in the methods.

The variant interference complexes were expressed and purified as described for the wild-typeTarget RNA and DNA cleavage by *Mtb* Csm The CRISPR array used in heterologous production of the Csm effector complex contained two copies of the same artificial spacer to minimise heterogeneity. We first evaluated the ability of Csm to cleave target RNA that was complementary to the crRNA. Incubation of Csm effector complex with 5’-radiolabeled target RNA (Figure 2, S2) in the presence of Mg^2+^ resulted in five cleavage products (B1 – B5) with the characteristic 6 nt spacing, consistent with other type III effectors (29-32). Non-complementary RNA was not cleaved by the effector complex (Figure S2A); however, anti-tag RNA with full-length complementarity to the crRNA (including 5’-handle), that mimics antisense RNA derived from the CRISPR locus, produced the same cleavage pattern as observed for target RNA (Figure S2B). A variant Csm complex with a mutation targeted to the cyclase domain (Cy) was indistinguishable from the wild-type protein for RNA cleavage. However, a variant (C3) with the Csm3 (Cas7) D35A mutation was not competent for target RNA cleavage, as expected (Figure 2) (30). We also investigated DNA cleavage by the *Mtb* Csm effector complex. We observed weak cleavage of ssDNA substrates by the wild-type complex; however, a variant with a mutation in the HD nuclease domain was still capable of degrading ssDNA, whilst the Csm3 D35A variant showed no DNase activity (Figure S3). No detectable hydrolysis was observed for DNA bubbles or dsDNA and the observed DNase activity was not dependent on the presence of target RNA. Together, these data suggest there is no functional HD nuclease activity in the recombinant *Mtb* Csm complex *in vitro*.

**Figure 2.**
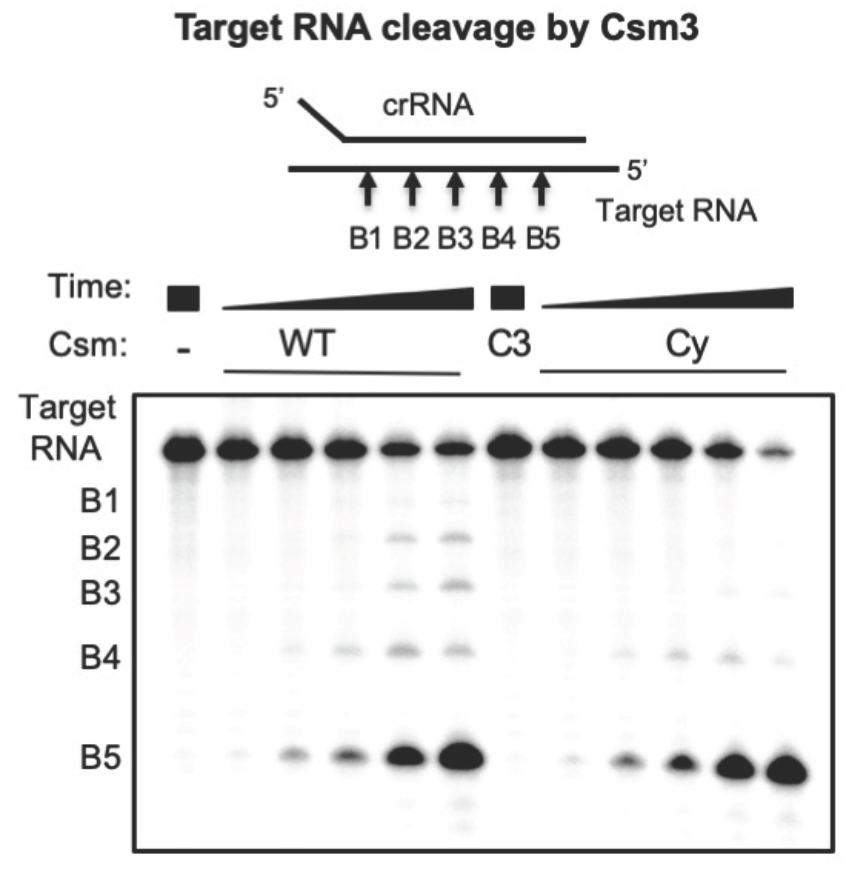
Target RNA cleavage by the Csm complex. Wild-type (WT), Csm3 D35A variant (C3), or cyclase variant (Cy) Csm were incubated with 5’-^32^P-target RNA under single-turnover conditions for 0.5, 2, 5, 30, 90 min and analysed by denaturing PAGE.

### Cyclic oligoadenylate synthesis by the Csm complex

Most type III CRISPR systems are closely associated with a gene encoding a Csm6 or Csx1 enzyme, which is activated by cOA generated by the Cas10 cyclase domain on target RNA binding (11,12) and required for immunity (17,33,34). To test for cOA production, we incubated the Csm interference complex with cold target RNA and ATP and analysed the products by LC-MS (Figure 3A). *Mtb* Csm produced cA_3_, cA_4_, cA_5_, cA_6_ in decreasing order of abundance. The identity of cA_4_ and cA_6_ was further confirmed by comparison to synt hetic standards (data not shown). We also detected substantial amounts of the corresponding linear OA triphosphates that are likely to be intermediates of cOA biosynthesis. When the same analysis was performed on metabolites produced by the Csm Csm3 D35A complex, a significant shift in cOA profile towards larger ring sizes was observed.

**Figure 3.**
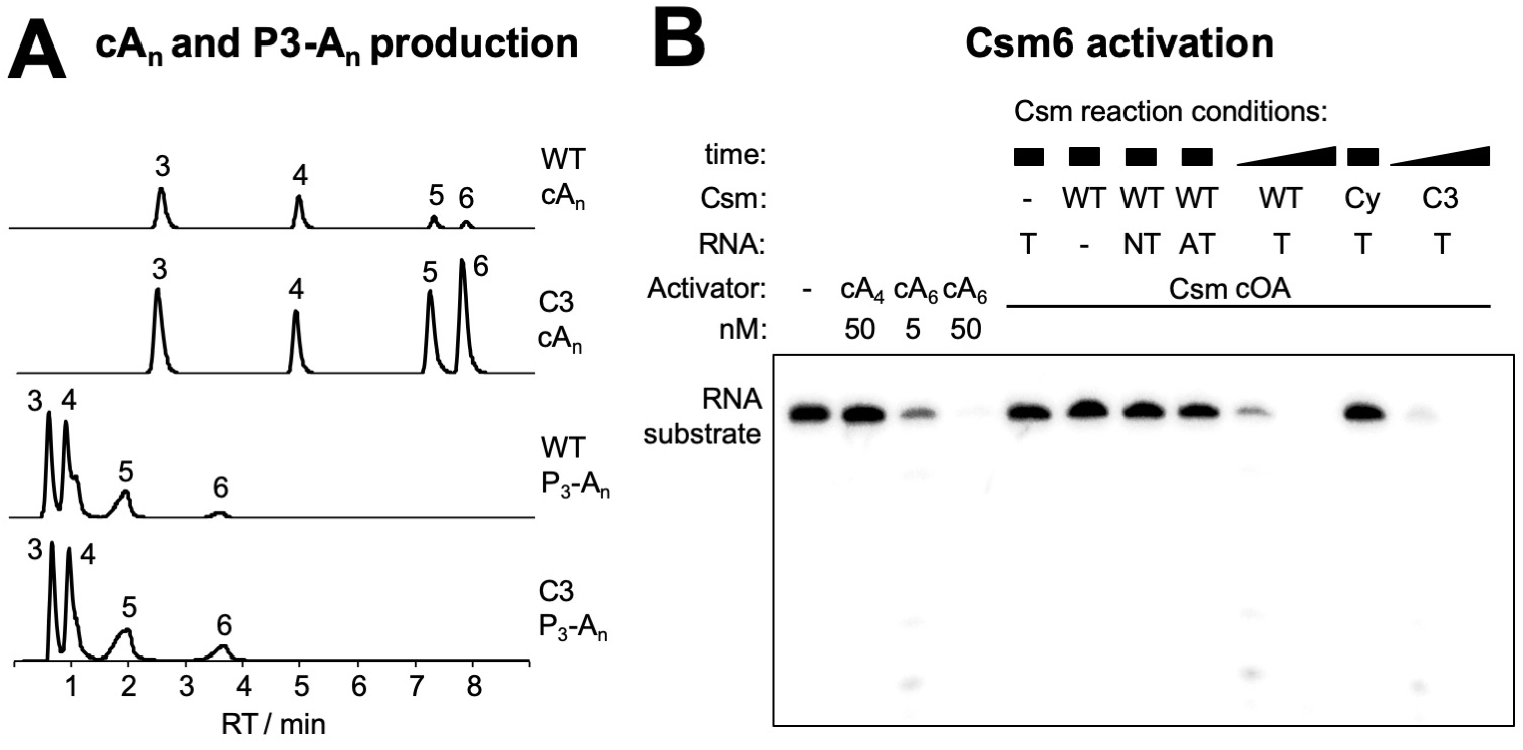
cOA production and Csm6 activation (**A**) Extracted ion chromatograms of oligoadenylates produced by wild type (WT) and Csm3 D35A variant (C3) Csm complex. Identical *y*-scaling throughout. The identity of the cyclic OAs (cA_n_) and respective 5’-triphosphates (P_3_-A_n_) is indicated by the number of AMP subunits. Production of cOAs, cA_5_ and cA_6_ in particular, was significantly increased for the Csm3 D35A (C3) Csm complex. (**B**) Csm6 ribonuclease activation by cyclic oligoadenylates. *Mtb* Csm6 (125 nM) was incubated with 200 nM 5’-[^32^P]-substrate RNA at 37 °C for 30 min in the presence of synthetic oligoadenylates (cA_4_, cA_6_) or a 10-fold dilution of Csm-derived cOAs. Csm wild-type (WT, 800 nM), cyclase variant (Cy, 800 nM), or Csm3 D35A (C3, 200 nM) was incubated with 1 mM ATP and 200 nM target (T), non-target (NT), or anti-tag (AT) RNA for 0.5, 2 (WT, C3), or 90 min (all others) at 30 °C. Only cA_6_ activated Csm6. For WT and C3 Csm, sufficient cA_6_ was produced within 0.5 min to switch on Csm6 ribonuclease activity; whereas no detectable amount of cA_6_ was produced under all other conditions even after prolonged reaction time.

### Activation of M. tuberculosis Csm6 requires cA_6_

The cOA-dependent ribonucleases studied to date are activated by either cA_6_ or cA_4_ (11-13). To determine which cOA species is required for activation of *Mtb* Csm6, we incubated the recombinant protein with synthetic cA_4_ and cA_6_, along with a radioactively labelled substrate RNA (Figure 3B). cA_4_ at 50 nM did not activate the ribonuclease activity, but in contrast cA_6_ showed strong activation at 5 nM and supported complete degradation of substrate RNA at 50 nM. This confirms cA_6_ as the relevant activator of *Mtb* Csm6. To investigate the activation of Csm6 by the Csm complex, we set up an assay where RNA binding and cOA synthesis by Csm are coupled to activation of Csm6 (13). In the presence of wild-type Csm and target RNA, strong activation of Csm6 was observed, consistent with the generation of cA_6_ by the Csm complex. Non-target (NT) control DNA did not activate Csm6, and neither did an RNA matching the 5’-handle of the crRNA (anti-tag, AT). The variant Csm with a mutation that inactivates the cyclase domain (Cy), as expected, did not activate Csm6, whereas the variant deficient in target RNA cleavage (C3) was, as expected, proficient in Csm6 activation, confirming previous findings for a variety of type III systems (11,13).

### In vivo activity of the Csm complex

We next set out to test if the Csm complex was active *in vivo*. A recent study of plasmid transformation in *M. tuberculosis* yielded 3 to 10-fold less transformants when the introduced plasmid carried one of the spacer sequences of the native CRISPR array, suggesting that the *Mtb* CRISPR system is functional in its native host (23). We tested the capability of Csm to confer immunity against incoming plasmids in the heterologous host *E. coli*. The genes for the Csm interference complex were assembled together with *csm6* (pCsm1-6, Table S1). pCsm1-6 was co-transformed with pCRISPR, harbouring the CRISPR array with *cas6* and two identical spacers targeting the pUC19 multiple cloning site (MCS) portion of the *lacZ*α gene. We constructed plasmid pRAT, which combines the origin of replication of pRSF-Duet-1 (Novagen), compatible with both Csm/CRISPR constructs, with tetracycline resistance and an arabinose-inducible MCS, and introduced pUC19 *lacZα*, containing the target sequence, into pRAT to give pRAT-Target. Plasmid maps are shown in Figure S4.

An *E. coli* C43 strain harbouring the complete *Mtb* Csm interference module, was transformed with 100 ng target plasmid pRAT-Target or with pRAT as negative control. Serial dilutions of the transformation mixture were applied to plates with three different conditions (Figure 4A): selecting for pCsm1-6 and pCRISPR only, additionally selecting for the incoming plasmid without induction of target expression, and finally selecting for the incoming plasmid with induction of target expression. The first condition provided the number of recipient cells and served to establish that differences in transformation efficiency were not due to different cell concentrations. The other two conditions were designed to establish if target transcription was essential for plasmid immunity. There was no significant difference in number of recipients. The number of transformants on uninduced plates also showed no significance difference between the control and target plasmid. When target expression was induced, however, approximately 100 times fewer transformants were observed for the target plasmid relative to control or uninduced conditions (Figure 4A). This demonstrates that the Csm-mediated immunity is functional in the heterologous host, and that immunity is dependent on target transcription as observed in other type III systems (35).

**Figure 4.**
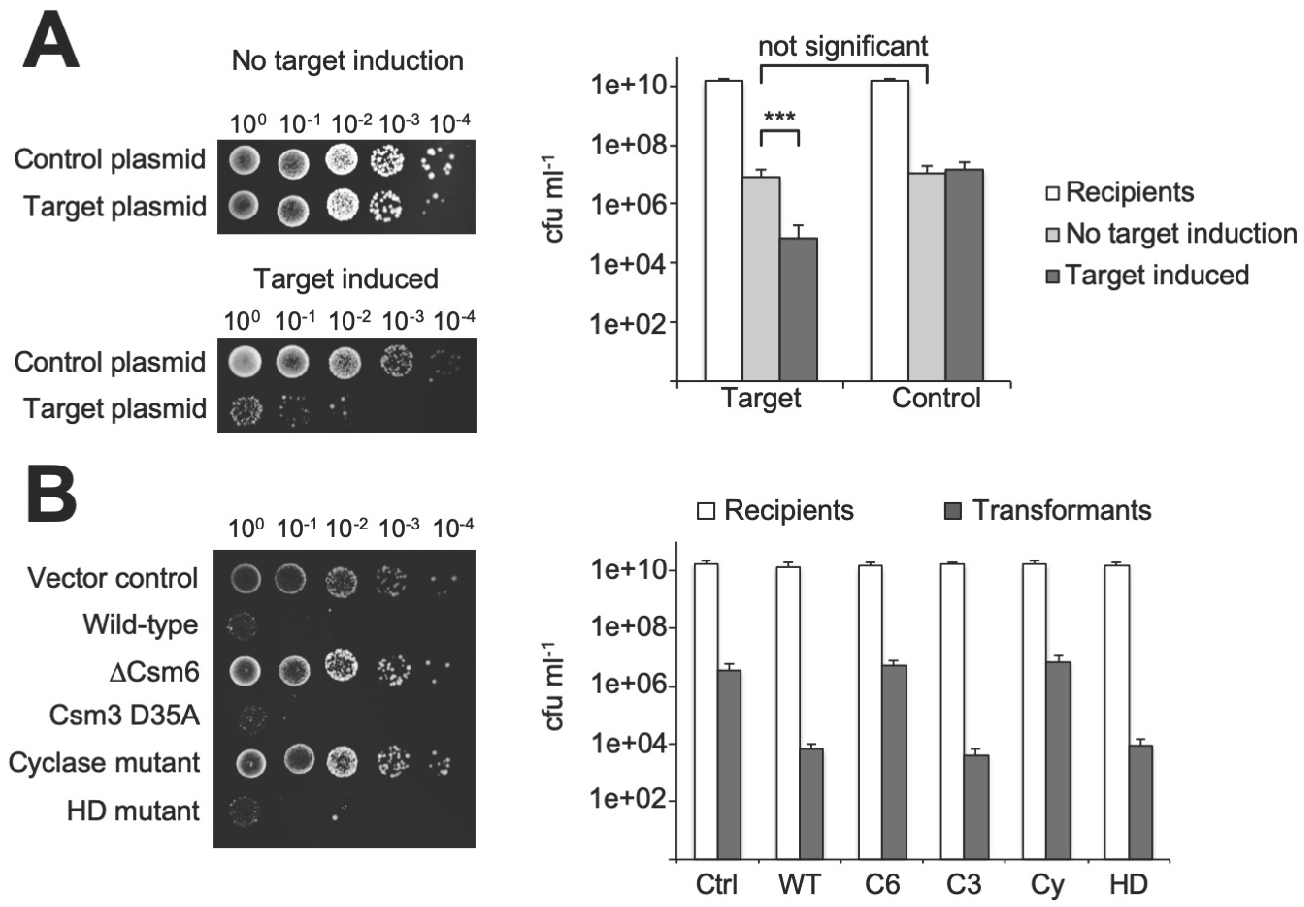
Plasmid immunity assays of *Mtb* Csm in heterologous host *E. coli*. (**A**) C43 harbouring pCsm1-6 and pCRISPR are transformed with pRAT (control plasmid) or with pRAT-target (plasmid with pUC19 MCS target). Target transcription was arabinose-inducible. In the absence of arabinose, the transformation efficiency was independent of the presence or absence of target; whereas when target transcription was induced, a 2-log reduction of transformation efficiency was observed for the target plasmid compared to the control plasmid. *** p-value (Welch two sample test, two tailed) < 1e-05; significance threshold at < 0.05. (**B**) C43 harbouring pCsm1-6 or indicated mutants and pCRISPR_TetR (target tetracycline resistance gene, constitutively expressed) were transformed with pRAT. Vector control (Ctrl): backbone vectors without inserts; wild-type Csm (WT), Csm1-5 only (ΔCsm6, C6), Csm3 D35A (C3), Csm1 (Cas10) D630A/D631A (cyclase mutant, Cy), Cas10 H18A/D19A (HD mutant, HD).

Thus far, the observed plasmid immunity was less efficient by one to three orders of magnitude compared to those in other type III systems (5,19,33,35,36). To test whether targeting a gene essential for plasmid maintenance would provide a higher degree of immunity than when targeting a non-essential one as before, we constructed a new CRISPR array with spacers targeting the tetracyline resistance gene of pRAT. The new array together with Cas6 was cloned into pCDFDuet-1 to give pCRISPR_TetR and co-transformed with the complete *Mtb* Csm module (pCsm1-6) into *E. coli* C43 as before. Constructs where *csm6* had been omitted (pCsm1-5_ ΔCsm6) or that carried inactivating mutations in the Csm3 subunit (pCsm1-6_ C3), the cyclase (pCsm1-6_ Cy) or the HD domain (pCsm1-6_ HD) were also constructed and introduced into *E. coli* together with pCRISPR_TetR. As a control, empty backbone (pCsm_Control) was co-transformed with pCDF-Duet-1. Cells were transformed with pRAT, and a 10-fold serial dilution was plated selecting for recipients (selection for pCsm and pCRISPR_TetR only) and for transformants (additional selection for incoming plasmid and induction of CRISPR-Cas genes).

We observed 3 orders of magnitude fewer transformants for the full wild-type Csm module (including Csm6) compared to the empty vector control (Figure 4B). As expected, the Csm3 active site variant (C3) was as active as wild type Csm; however, so was the Csm HD variant. No plasmid immunity was observed for the Csm cyclase domain variant (Cy) or when Csm6 was not present. This is strong indication that cOA induced Csm6 ribonuclease activity is the driving force behind plasmid immunity for this system. It also demonstrates that target RNA cleavage is not required for immunity, nor is DNase activity, at least not in a heterologous host such as *E. coli*.

### Reprogramming the Mtb Csm system as a cA_4_-mediated defence

The observation that the *Mtb* Csm complex generates a range of cOA species from cA_3_-cA_6_ is consistent with previous findings for type III systems (11-13). Thus, the cOA species relevant for downstream immunity should be defined by the CARF-family effector protein(s) present in the organism. For *Mtb*, the native Csm6 enzyme is activated by cA_6_. We sought to determine whether we could reprogramme the *Mtb* CRISPR system as a cA_4_-mediated defence, by replacing Csm6 with an alternative effector protein. A candidate Csx1 protein from the gammaproteobacterium *Thioalkalivibrio sulfidiphilus* (37) was selected and expressed in *E. coli* from a synthetic gene. The activity of the purified recombinant protein (TsuCsx1) was tested against a labelled RNA substrate in the absence and presence of cOA species (Figure 5A). TsuCsx1 was strongly activated by synthetic cA_4_ but not by cA_6_, demonstrating that this enzyme is cA_4_ specific. Strong activation was also observed when the enzyme was incubated with cOA generated by the *Mtb* Csm complex, consistent with the observation that cA_4_ is a significant component of this mixture (Figure 3A).

**Figure 5.**
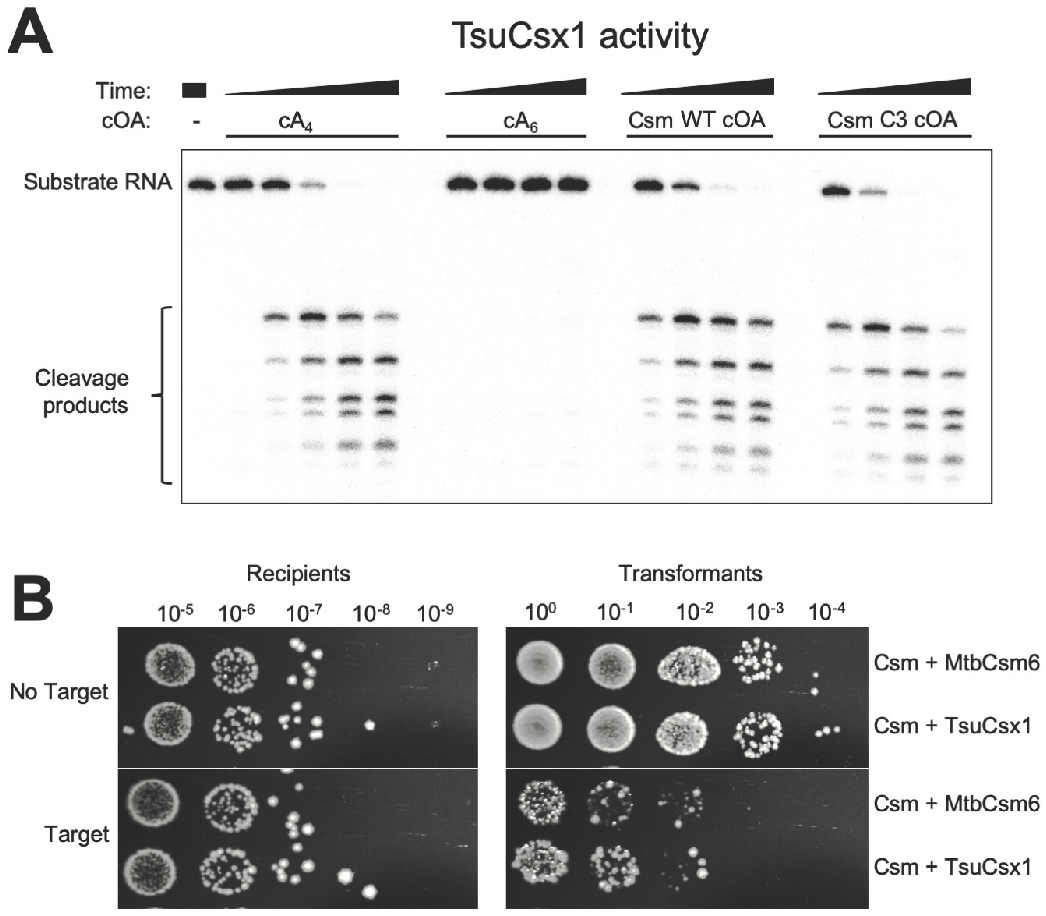
Re-programming the *Mtb* Csm system for cA_4_-responsive immunity. (**A**) *In vitro* activity of TsuCsx1. The reaction contained 0.5 µM TsuCsx1 dimer, 100 nM 5’-^32^P-labeled RNA A1, 20 mM Tris, 150 mM NaCl, 1 mM DTT, pH 7.5 and was conducted at 35 °C for 1, 5, 15, 30 min. Activators were added as indicated; Csm-derived cOAs were from a 2 h reaction, otherwise as described for Figure 3. TsuCsx1 was activated by cA_4_ but not cA_6_, and Csm-derived cOAs are able to induce Csx1 ribonuclease activity *in vitro*. (**B**) Plasmid immunity assay using pUC19 *lacZα*-targeting Csm effector complex in *E. coli* C43 (pCsm1-5_Csm6/tsuCsx1 and pCRISPR, Figure S4). The ribonuclease was either the cognate Csm6 or TsuCsx1. Cells were transformed with pRAT (control plasmid) or pRAT-Target (target plasmid) and 10-fold serial dilutions were plated on selective plates containing arabinose for induction of target transcription. TsuCsx1 confers the same level of plasmid immunity as the cognate *Mtb* Csm6.

We proceeded to add the gene encoding TsuCsx1 to the *Mtb csm1-5* genes (pCsm1-5_Csx1). As before, *E. coli* C43 harbouring pCsm1-5_tsuCsx1 and pCRISPR, were transformed with the target plasmid (pRAT-Target) and pRAT as the control lacking the target sequence. Ten-fold serial dilutions were applied to selective, inducing plates to determine the number of transformants; the cell count for recipients was carried out as before. We observed similar levels of plasmid immunity for the cognate *Mtb* Csm6 and the heterologous TsuCsx1 systems (Figure 5B), demonstrating that the downstream effector proteins determine which cyclic oligoadenylate species is active for immunity, and that the Csm interference complex is not specific for a given ribonuclease or *vice versa*

## DISCUSSION

### The type III-A (Csm) CRISPR system of M. tuberculosis provides immunity via cOA signalling

*M. tuberculosis* harbours the archetypal type III-A (“Mtube”) CRISPR system, and the variability of CRISPR loci in mycobacteria (spoligotyping) has been utilized to classify isolates for over 20 years. Despite this, little has been reported on the characterisation of the *Mtb* CRISPR system, particularly *in vitro*. Initial work on reconstituting the *Mtb* Csm complex in *E. coli* encountered problems due to poor solubility of the Csm2 (Cas11) subunit. We surmounted this hurdle by swapping the problematic *Mtb csm2* gene with that from the closely related species *M. cannetti*, allowing expression of *Mtb* Csm complex that is functional both *in vitro* and *in vivo*. Such an approach may find utility in other circumstances when studying multi-protein complexes.

Having succeeded in constructing an active *Mtb* Csm complex, we could dissect the contributions of the various enzymatic activities of this complex to CRISPR-based immunity. We could detect no strong evidence for target RNA dependent DNA cleavage by the HD nuclease domain of the *Mtb* Csm complex (Figure S3). Furthermore, targeted mutation of the HD nuclease active site did not affect the ability of the *Mtb* CRISPR system to provide immunity against plasmid transformation in *E. coli*, and immunity was lost when the Cyclase active site was inactivated and the HD nuclease unaffected (Figure 4). Target RNA dependent DNA cleavage by the HD nuclease domain of Cas10 has been demonstrated *in vitro* for the type III-A complexes of *S. epidermidis* (38), *S. thermophilus* (7) and *Thermus thermophilus* (39), and type III-B complexes of *Thermatoga maritima* (6), *Pyrococcus furiosus* (5) and *S. islandicus* (9). On the other hand, not all type III CRISPR effectors support robust DNase activity *in vitro* (32), and not all type III systems have an intact HD nuclease domain (3). HD nuclease activity is often not required for plasmid immunity *in vivo*, although the cyclase activity of Cas10 and the presence of a cOA effector nuclease clearly is (9,17-19). Recent analysis of plasmid clearance by the type III-A system of *S. epidermidis* revealed that highly transcribed plasmid targets were cleared efficiently by DNA targeting, whilst low levels of transcription resulted in a requirement for RNA cleavage by the Csm6 nuclease. This suggests that these two aspects of type III defence systems can work in tandem, and assume differing importance for immunity, depending on the biology of the invading genetic element (36). Although we see no target-activated role for the HD nuclease domain of *Mtb* Csm, either *in vitro* or in *E. coli*, we cannot rule out a role *in vivo* in the homologous host. It is possible that the recombinant system generated here has resulted in deactivation of the HD nuclease, for example, or that a missing component is crucial for full function in *M. tuberculosis*.

Cyclic oligoadenylate synthesis has been studied for a variety of type III CRISPR systems. One emerging paradigm is that a range of cOA molecules, from cA_3_ to cA_6_, are synthesised – at least for recombinant systems *in vitro*. We observed a predominance of the smaller species for wild-type *Mtb* Csm (Figure 3A), consistent with observations for the type III-A complex from *S. thermophilus* (11). Notably, when target RNA cleavage was abolished by introduction of a D35A mutation in the Csm3 subunit, the levels and distribution of cOA species changed dramatically. A significant increase in the amounts of the larger cA_6_ and cA_5_ species was observed, with cA_6_ coming to predominate. This reflects the finding that target RNA cleavage by the Csm3 subunit is the event that deactivates the cyclase domain (13,40). However, even with target RNA cleavage abolished, a range of cOA species are synthesised. This leads to the question: is this a feature or a bug? Or in other words, has the ability to generate a range of cOA species been selected for by evolution, or just not selected against?

### Ancillary defence enzymes are replaceable parts for Type III CRISPR systems

A number of Csx1/Csm6 family proteins have now been studied structurally and/or biochemically (Table 1). As already noted, proteins in this family have an N-terminal CARF domain and a C-terminal HEPN domain. They tend to be named as “Csm6” proteins if their genes are closely associated with those encoding a type III-A/Csm CRISPR effector complex, and Csx1 proteins if they are not, but it is not clear that this division relates to any intrinsic property or sequence of family members. The enzymes from the Archaeal, Deinococcus-Thermus and Proteobacterial phlya that have been studied to date all utilise cA_4_ as an activator (Table 1), with examples of cA_6_-dependent enzymes only characterised so far in the Firmicutes and Actinobacteria. As cA_4_ and cA_6_ have quite significantly different structures and sizes, this is likely to be reflected in the structures of the corresponding CARF domains that bind these ligands. To date however, there is no structural information available for any cA_6_-binding CARF domain – this is a priority for future studies.

**Table 1.** Comparison of characterised Csx1/Csm6 family proteins.

| Species | Phylum | Gene | CRISPR subtype | PDB | cOA activator | Reference |
| --- | --- | --- | --- | --- | --- | --- |
| *S. solfataricus* | Crenarchaeota | Sso1389 | III-D | 2I71 | cA<sub>4</sub> | (13) |
| *S. islandicus* | Crenarchaeota | SiRe_0884 | III-B |  | cA<sub>4</sub> | (47) |
| *P. furiosus* | Euryarchaeota | PF1127 | III-B | 4EO G | ? | (48) |
| *M. thermautotrophicus* | Euryarchaeota | MTH1076 | III-A |  | cA<sub>4</sub> | (32) |
| *T. thermophilus* | Deinococcus-Thermus | TTHB152 | III-A | 5FSH | cA<sub>4</sub> | (12) |
| *T. thermophilus* | Deinococcus-Thermus | TTHB144 | III-A |  | cA<sub>4</sub> | (49) |
| *T. sulfidophilus* | Proteobacteria | Tgr7_1803 | III-B |  | cA<sub>4</sub> | This work |
| *S. thermophilus* | Firmicutes | StCsm6 | III-A |  | cA<sub>6</sub> | (11) |
| *E. italicus* | Firmicutes | EiCsm6 | III-A |  | cA<sub>6</sub> | (23) |
| *M. tuberculosis* | Actinobacteria | Rv2818c | III-A |  | cA<sub>6</sub> | This work |

\* Activator has not been determined for PF1127, but predicted to be cA<sub>4</sub>.

Based on the data collated in Table 1, it is already apparent that the cOA activator utilized by any particular type III system does not correlate with the CRISPR subtype. Although only type III-A systems are known to utilize cA_6_, there are well-studied examples (such as *T. thermophilus* and *M. thermautotrophicus*) of type III-A systems with ancillary ribonucleases activated by cA_4_. All type III systems studied to date have the capability of making a range of cyclic oligoadenylates from cA_3_-cA_6_ (11-13,40). As we have shown here, the immunity provided by a type III system can be readily converted from a cA_6_ to a cA_4_-based mechanism, simply by replacing the ancillary enzyme. This suggests there is no direct cross-talk between the type III complex and the ribonuclease, other than via the cOA activator. Given the extent of gene loss and gain in CRISPR loci, the capture of a new CARF-family effector protein might be sufficient to potentiate a switch from one cOA signalling molecule to another. It is equally possible that distinct effector proteins, activated by cA_4_ and cA_6_, could operate in parallel in a single host species. This might conceivably provide enhanced levels of immunity.

### Cyclic and linear oligoadenylate signalling – more to discover?

Another pressing question for type III CRISPR systems concerns the role (if any) for cA_3_ molecules, which are often the most abundant species synthesised, at least *in vitro* (11,12). Given the lack of 2-fold symmetry for cA_3_, recognition of this molecule may be accomplished by a sensing domain quite distinct from the dimeric CARF domain found in many CRISPR-associated proteins (13). Recent discoveries of novel cyclic trinucleotides synthesised by bacterial cGAS-like enzymes and detected by vertebrate innate immune surveillance proteins (41,42) make this question pertinent. This is particularly true for organisms such as *M. tuberculosis*, which has an intracellular lifestyle in vertebrate hosts with which it participates in a two-way communication via cyclic nucleotides (43,44). Furthermore, the significant quantities of nanoRNAs (eg A_3_-triphosphate) synthesised by the Csm complex should not be discounted as dead-end side products. NanoRNAs have been shown capable of priming transcription initiation in bacteria (45). Recently, the CnpB (Rv2837c) protein, which can degrade nanoRNAs, controls the transcription of the *csm* genes and CRISPR RNA in *M. tuberculosis* (46). Deletion of the *cnpB* gene leads to massive transcriptional up-regulation of the *csm* genes, which is consistent with a model whereby nanoRNAs generated by the Csm cyclase domain play a role in sculpting the transcriptional response to phage infection in cells with a type III CRISPR system by activating defence gene expression. Clearly, there are still many avenues to explore.

## ACKNOWLEDGEMENTS

Thanks to Christophe Rouillon, Marialena Varympopioti & Chloe Jones for helpful discussions.

## SUPPLEMENTARY INFORMATION

**Figure S1.**
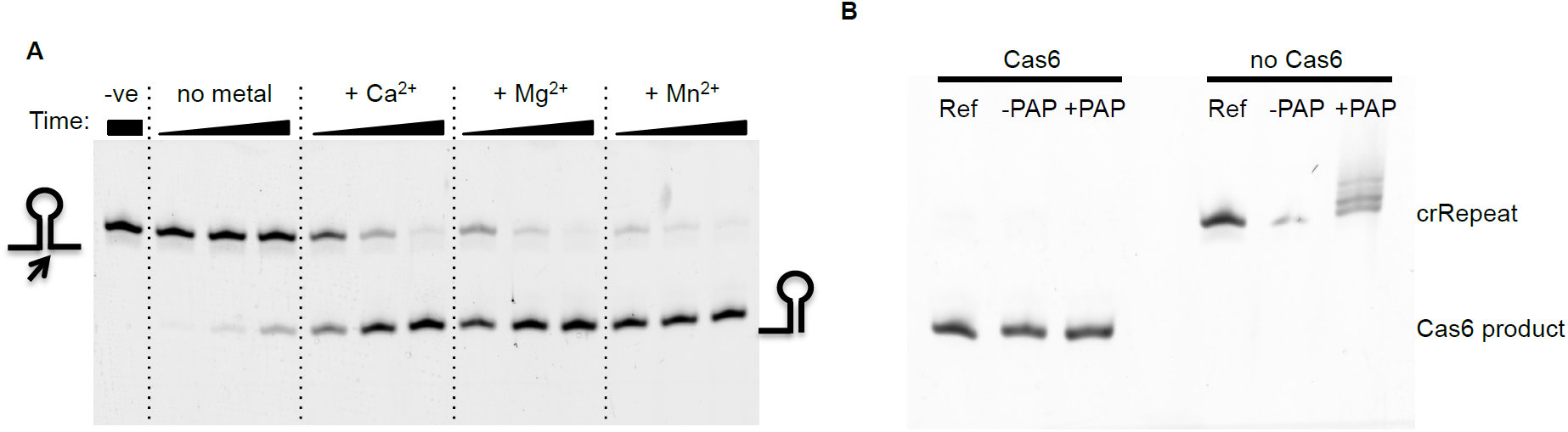
The Cas6 ribonuclease cleaves the *Mtb* CRISPR repeat sequence to generate crRNA. (**A**) Cas6 (0.5 µM) was incubated with 50 nM 5’-FAM CRISPR repeat RNA at 37 °C for 5, 15, 45 min in 20 mM Tris, 100 mM potassium glutamate, pH 7.5 in the absence or presence of 5 mM divalent metal ions as indicated. Reactions were stopped by phenol-chloroform extraction. (**B**) Cas6 cleavage leaves a 3’-(cyclic) phosphate group. CRISPR repeat RNA (crRepeat, 5’-FAM labeled, 400 nM) was digested with 2 µM Cas6 for 1 h in the presence of Mg^2+^ using the same reaction conditions as before. Phenol-chloroform followed by chloroform extraction provided the substrate for the *E. coli* Poly(A) polymerase (PAP, New England Biolabs) reaction. Polyadenylation was performed according to the manufacturer’s instructions. In a parallel experiment, Cas6 was omitted. The CRISPR repeat RNA but not the Cas6 product can be 3’-polyadenylated by PAP. This suggests that the reaction product has a cyclic 2’,3’-phosphate, as observed for other Cas6 enzymes. This observation, together with the observation that calcium supports enhanced cleavage of the CRISPR repeat, suggests that the metal ion does not participate directly in catalysis but rather plays a role in stabilisation of the RNA substrate or RNA:protein complex.

**Figure S2.**
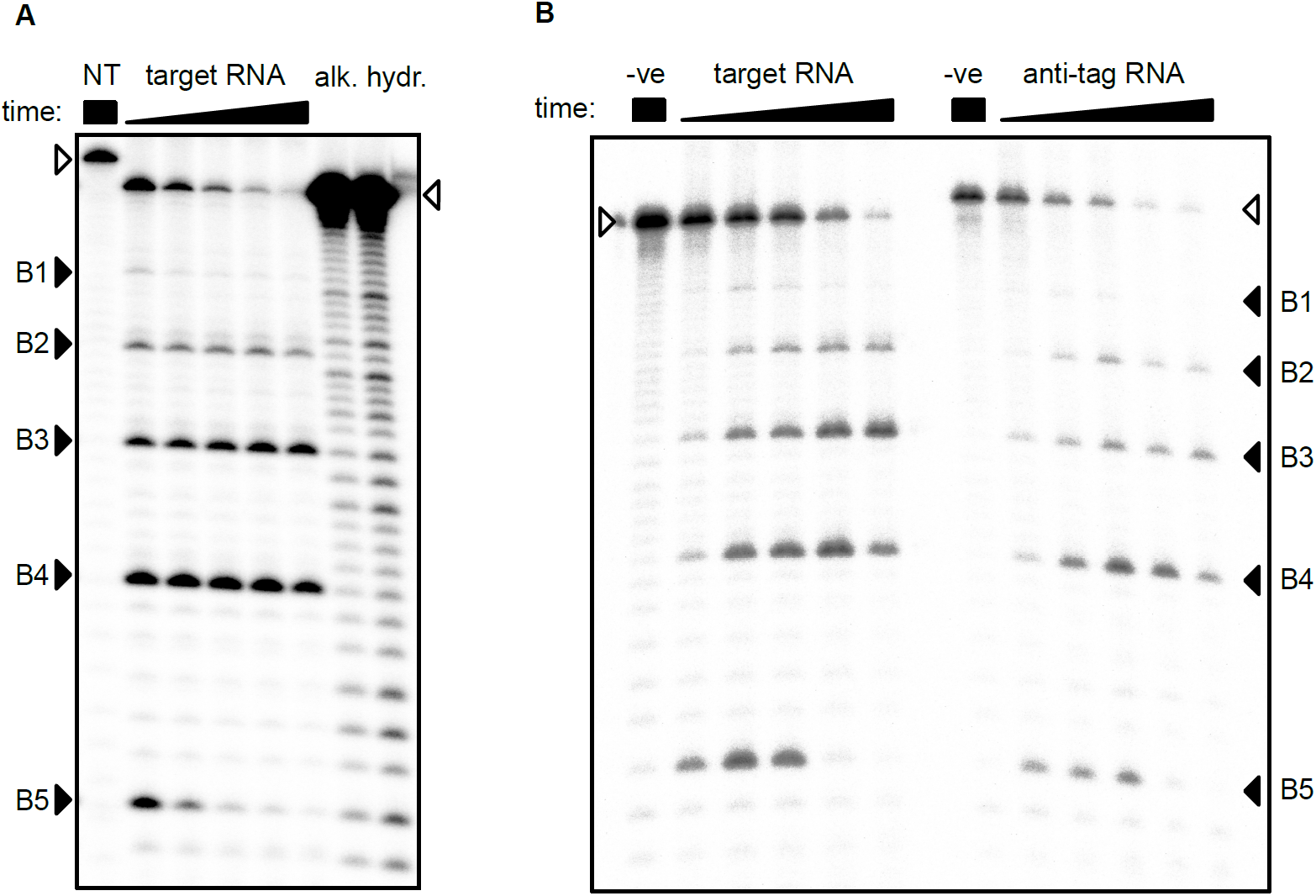
RNA backbone cleavage by Csm interference complex. **(A**) 5’-^32^P-labeled target RNA or non-target (NT) RNA were treated with 0.8 µM Csm effector complex for 5, 15, 30, 60, 120 min. Reactions were analysed by denaturing PAGE alongside alkaline hydrolysis ladders prepared from target RNA. (**B**) Target RNA and anti-tag RNA (full-length complementarity to crRNA including repeat-derived 5’-handle) are both cleaved by Csm to give identical products. Reaction time: 0.5, 2, 5, 30, 100 min; substrate RNA is indicated by unfilled triangles; the five cleavage sites (B1 – B5) with the characteristic 6 nt spacing are indicated by filled triangles. Target RNA with a 4 nt truncation at the 3’-end was used, leading to a different distribution of cleavage products compared to Figure 2.

**Figure S3.**
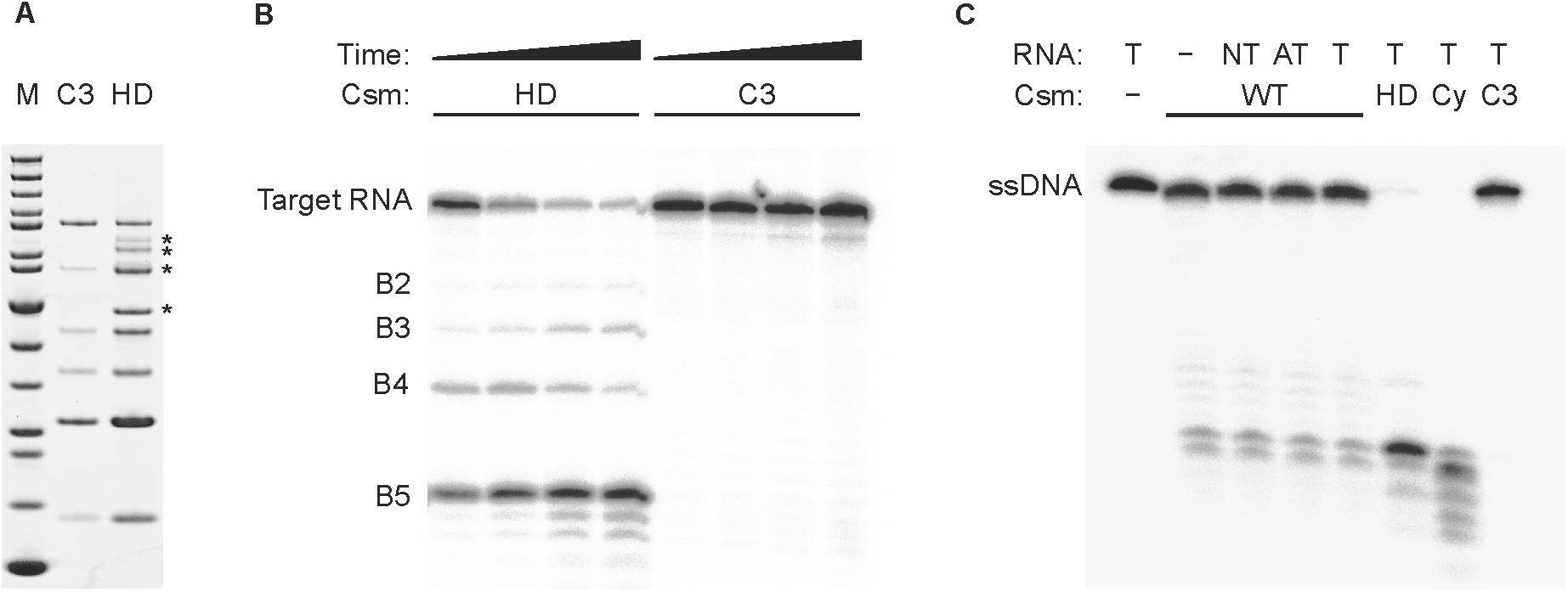
In vitro analysis of Csm HD mutant. (**A**) SDS-PAGE of purified Csm3 D35A (C3) and Cas10 H18A/D19A (HD) variant interference complexes; *: contaminants; M: PageRuler Unstained (Fisher Scientific). (**B**) Target RNA backbone cleavage by Csm3 D35A (C3) and Cas10 H18A/D19A (HD) variant interference complexes; 5’-radiolabeled target RNA was incubated with HD or C3 in the presence of Mg^2+^ for 5, 10, 30, 60 min at 30 °C; as expected the characteristic cleavage products B2 – B5 are produced by HD but not the C3 variant. (**C**) The ssDNase activity of Csm interference complex is not dependent on RNA substrate or active site mutations; 5’-radiolabeled ssDNA was incubated with Csm wild type (WT), HD, Cas10 D630A/D631A (Cy), or Csm3 D35A (C3) in the presence of Mg^2+^ for 90 min at 30 °C; cold RNA was added as indicated. Mn^2+^ and Co^2+^ also supported the observed activity, Zn^2+^ less so, and Cu^2+^ did not stimulate DNase activity (data not shown). T: target RNA, NT: non-target RNA, AT: anti-tag RNA.

**Figure S4.**
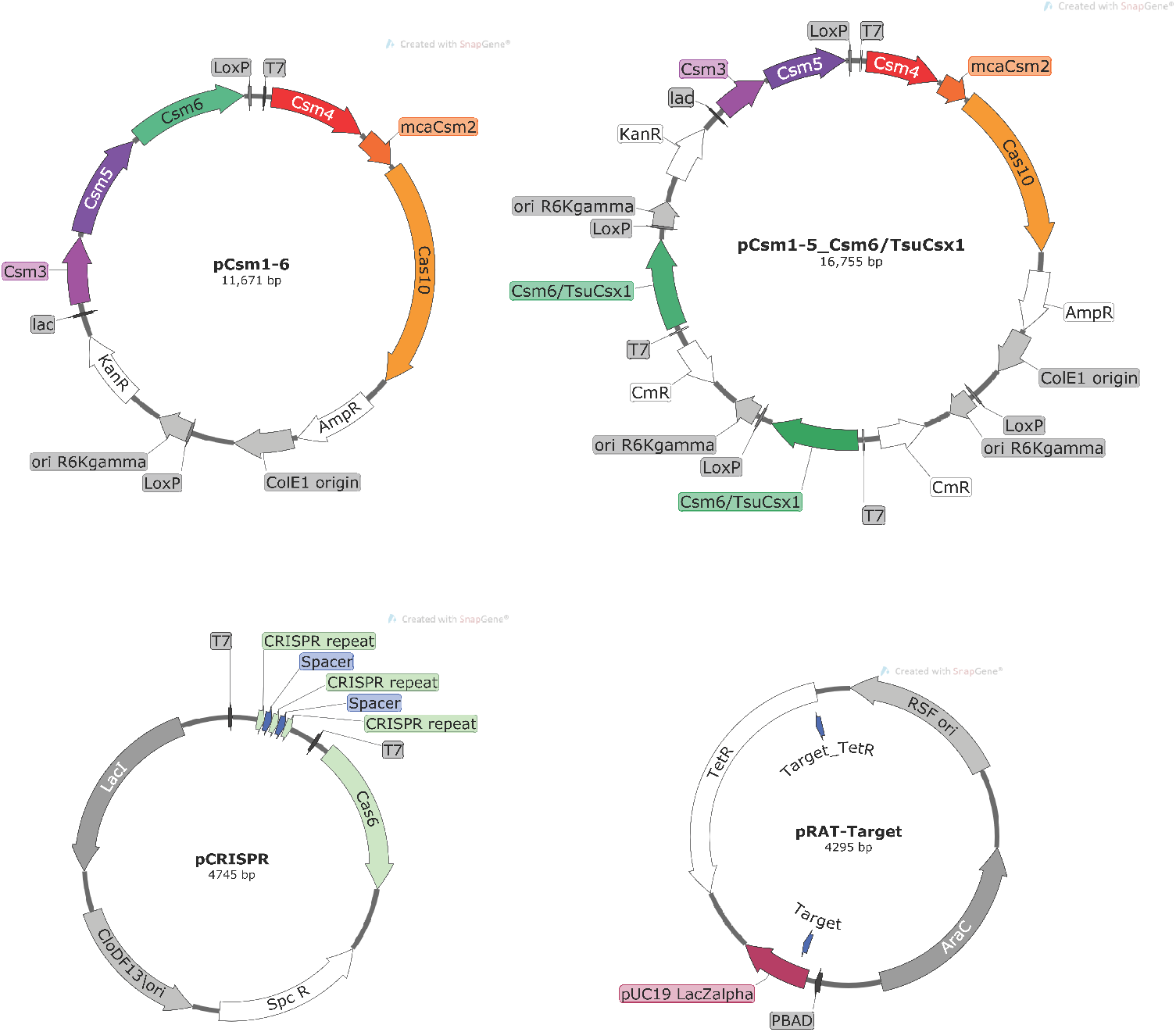
Selected plasmid maps for constructs used in plasmid immunity assays. In pRAT-Target, the blue arrows indicate the positions of the match to crRNA targeting the pUC19 LacZ□ MCS (Target) or the tetracycline resistance gene (Target_TetR).

**Table S1:** Plasmids constructed in this study.

| Name | Genes | Resistance | Notes | Use |
| --- | --- | --- | --- | --- |
| pCsm1-5 | *csm1*, *csm2*, *csm3*,<br>*csm4*, *csm5*,<br>*Mca_csm2* | Amp,<br>Kan, Cm | MultiColi™ pACE, pDK, pDC recombination; ColE1 ori; pT7 (*csm1*, 2, 4 on pACE), pLac (*csm3*, 5 on pDK), pT7 (*Mca_csm2* on pDC), TEV-cleavable N-His tag on Csm4 | Protein production and purification |
| pCsm1-5_C3 | *csm1*, *csm3* *D35A*,<br>*csm4*, *csm5*,<br>*Mca_csm2* | Amp, Kan | MultiColi™ pACE and pDK recombination; ColE1 ori; pT7 (*csm1*, 4, *Mca_csm2* on pACE), pLac (*csm3*, 5 on pDK), TEV-cleavable N-His tag on Csm4 | Protein production and purification |
| pCsm1-5_Cy | *csm1* *D630A*<br>*D631A*, *csm3*, *csm4*,<br>*csm5*, *Mca_csm2* | Amp, Kan | as pCsm1-5_C3 | Protein production and purification |
| pCsm1-6 | *csm1*, *csm3*, *csm4*,<br>*csm5*, *csm6*,<br>*Mca_csm2* | Amp, Kan | pLac (*csm3*, 5, 6), otherwise as pCsm1-5_C3 | Plasmid immunity assay |
| pCsm_Control | - | Amp, Kan | pACE and pDK recombination | Plasmid immunity assay; vector control |
| pCsm1-5_ΔCsm6 | *csm1*, *csm3*, *csm4*,<br>*csm5*, *Mca_csm2* | Amp, Kan | as pCsm1-5_C3 | Plasmid immunity assay |
| pCsm1-6_C3 | *csm1*, *csm3* *D35A*,<br>*csm4*, *csm5*, *csm6*,<br>*Mca_csm2* | Amp, Kan | as pCsm1-6 | Plasmid immunity assay |
| pCsm1-6_Cy | *csm1* *D630A*<br>*D631A*, *csm3*, *csm4*,<br>*csm5*, *csm6*,<br>*Mca_csm2* | Amp, Kan | as pCsm1-6 | Plasmid immunity assay |
| pCsm1-6_HD | *csm1* *H18A* *D19A*,<br>*csm3*, *csm4*, *csm5*,<br>*csm6*, *Mca_csm2* | Amp, Kan | as pCsm1-6 | Plasmid immunity assay |
| pCsm1-5_Csm6 | *csm1*, *csm3*, *csm4*,<br>*csm5*, *csm6*,<br>*Mca_csm2* | Amp,<br>Kan, Cm | pACE, pDK, pDC recombination; pT7 (*csm6* on pDC), otherwise as pCsm1-5_C3 | Plasmid immunity assay |
| pCsm1-5_tsuCsx1 | *csm1*, *csm3*, *csm4*,<br>*csm5*, *Mca_csm2*,<br>*Tsu_csx1* | Amp,<br>Kan, Cm | as pCsm1-5_Csm6 but *Tsu_csx1* instead of *csm6* | Plasmid immunity assay |
| pCRISPR | *cas6*, pUC MCS-targeting CRISPR array (3 repeats, 2 spacers) | Spc | pCDF-Duet-1 derivative; CRISPR in MCS-1, *cas6* in MCS-2 | crRNA biogenesis, *in vitro* and *in vivo* |
| pCRISPR_TetR | *cas6*, TetR-targeting CRISPR array (5 repeats, 4 spacers) | Spc | as pCRISPR | crRNA biogenesis, *in vivo* |
| pRAT | - | Tet | RSF ori; pBAD | Plasmid immunity assay; control plasmid for pUC MCS-targeting Csm, target plasmid for TetR-targeting Csm |
| pRAT-Target | pUC19 *lacZα* | Tet | as pRAT | Plasmid immunity assay; target plasmid for pUC MCS-targeting Csm |
| pCas6-CHis | *cas6* | Kan | pEHisTEV-derivative; C-terminal His-tag | Protein production and purification |
| pEHisTEV-Csm6 | *csm6* | Kan | pEHisTEV-derivative; TEV-cleavable N-His tag | Protein production and purification |
| pEHisTEV-TsuCsx1 | *Tsu_csx1* | Kan | pEHisTEV-derivative; TEV-cleavable N-His tag | Protein production and purification |

**Table S2.**
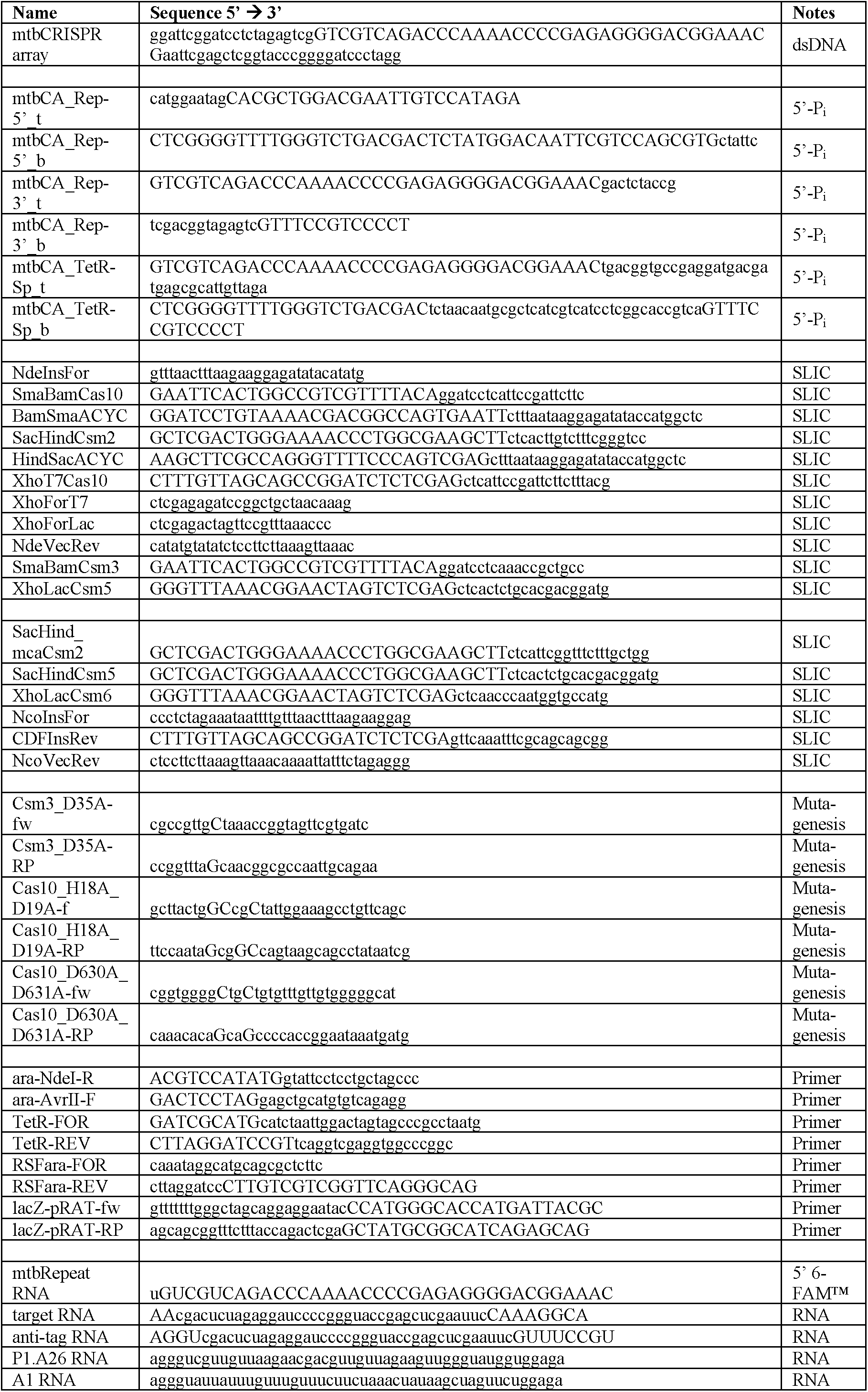
Oligonucleotides and primers used in this study.

